# Components of Executive Function Predict Regional Prefrontal Volumes

**DOI:** 10.1101/374009

**Authors:** Ryan A. Mace, Abigail B. Waters, Kayle S. Sawyer, Taylor Turrisi, David A. Gansler

## Abstract

**Objective:** Designed to measure a diversity of executive functioning (EF) through classical neuropsychological tests, the Delis-Kaplan Executive Function Scale (D-KEFS) allows for the investigation of the neural architecture of EF. We examined how the D-KEFS Tower, Verbal Fluency, Design Fluency, Color–Word Interference, and Trail Making Test tasks related to regional frontal lobe volumes, quantifying how components of EF were represented in disparate neural networks.

**Method:** Adults from the Nathan Kline Institute – Rockland Sample (NKI-RS), an open-access community study of brain development, with complete MRI (3T scanner) and D-KEFS data were selected for analysis (*N* = 478; ages 20-85). In a mixed-effects model predicting volume, D-KEFS task, D-KEFS score, region of interest (ROI; 13 frontal, 1 occipital control), were entered as fixed effects with intercepts for participants as random effects.

**Results:** “Unitary” EF (average of D-KEFS scores) was positively associated with superior frontal, rostral middle frontal, and lateral orbitofrontal volumes; a negative association was observed with frontal pole volume (| *z*-score slope | range = 0.040 to 0.051). “Diverse” EF skills (individual D-KEFS task scores) were differentially associated with two or three ROIs, respectively, but to a stronger extent (| *z*-score slope | range = 0.053 to 0.103).

**Conclusions:** The neural correlates found for the D-KEFS support the prefrontal modularity of EF at both the unitary (aspects of EF ability common to all tasks) and task (diverse EF skills) levels. The separation of task-general variance in neurocognition from task-specific variance can further evaluate neuropsychological tests as indices of brain integrity.

**Public Significance Statements:**

1. Our results support the relationship between larger lateral prefrontal cortex and greater executive function.
2. A composite of executive function performance, which was broadly associated with the prefrontal cortex, may be ideal for assessing diffuse frontal lobe damage (e.g., hypoxia).
3. Individual executive functions, which were more narrowly but strongly related to specific prefrontal regions, could be better for assessing the effects of localized brain injuries (e.g., tumor).

## Introduction

Executive function (EF) is a term that has been used by neuropsychologists for several decades. While its definition is not consistently applied, EF refers to cognitive processes that plan, coordinate, and regulate behavior and cognition (Diamond, 2013). These higher-order mental abilities include set-shifting, updating of working memory, and response inhibition (Miyake et al., 2000). The traditional definition of EF indicated that the prefrontal cortex was largely responsible for these skills (Luria, 1966; Teuber, 1972). The identification of brain-behavior relationships is a central aim of clinical and experimental neuropsychology. Such investigations are predicated on fundamental assumptions in the field: (1) the cerebral cortex possesses a high degree of specialization, (2) brain function can be inferred via a modularity approach to analyzing complex cognitive skills, and (3) brain damage can selectively disrupt subcomponents of a cognitive system (Cipolotti & Warrington, 1995). Investigating prefrontal cortex modularity of EF in non-patient samples are a prerequisite for understanding the nature of those systems under neuropathological circumstances. Advances in quantitative digital imaging support more refined research on EF and frontal brain regions guided by this theoretical framework (Bigler, 2013).

The nature of EF is a topic of considerable clinical and academic interest given its widespread role in everyday functioning (Diamond, 2013). EF deficits are a strong predictor of disability (Royall et al., 2007), general cognitive decline (Salthouse, Atkinson, & Berish, 2003), decreased mental health (Gansler, Suvak, Arean, & Alexopoulos, 2015), and increased mortality (Johnson, Lui, & Yaffe, 2007). Yet, there is significant debate on the conceptualization EF. For example, EF has been theorized as: the “top-down” inhibition of “bottom-up” processing (Aron, 2007); a multifaceted model of working memory (Baddeley & Hitch, 1974); cognitive control of the prefrontal cortex (Miller & Cohen, 2001), and being comprised of broad higher-order components (Lezak, 1995; Luria, 1966). The approach that the present work built upon is the EF model constructed by Miyake and Friedman, which examines the extent to which EF consists of separate “diverse” (i.e., distinct) components (shifting, updating, inhibition), and the extent to which it exists as a “unitary” construct (Miyake et al., 2000).

The Delis-Kaplan Executive Function Scale (D-KEFS) is a battery designed to measure EF (Delis, Kaplan, & Kramer, 2001) with normative data from a representative, large national sample (Homack, Lee, & Riccio, 2005). The D-KEFS, and the classical paradigms upon which its tasks are based, have been widely used in both clinical and research settings to assess executive function (Rabin, Barr, & Burton, 2005). One of the development aims of the D-KEFS was to take various classical measurement paradigms within clinical neuropsychology that were separately developed and co-norm the paradigms. The D-KEFS makes multiple EF tasks available (e.g. Trail Making Test, Color Word Interference, Design Fluency, Verbal Fluency, Tower) so that the clinician can tailor specific cognitive tests to the referral question and relevant clinical issue (Delis et al., 2001). Scores from these individual tasks may be interpreted as the “diverse” aspects of EF, as in the Miyake et al. (2000)model. The battery is an adequately reliable measure of EF (Delis, Kramer, Kaplan, & Holdnack, 2004) with discriminant validity in clinical populations across the lifespan (Heled, Hoofien, Margalit, Natovich, & Agranov, 2012; McDonald, Delis, Norman, Tecoma, & Iragui, 2005; Wodka et al., 2008).

Like many traditional neuropsychological tests of EF, the D-KEFS has been validated with lesion studies that show marked deficits for both individual aspects of EF (Strong, Tiesma, & Donders, 2011; Yochim, Baldo, Nelson, & Delis, 2007) and a common EF factor (Barbey et al., 2012). Systematic reviews of the neuroimaging literature show that among neuropsychiatric populations (Nowrangi, Lyketsos, Rao, & Munro, 2014) and healthy controls (Yuan & Raz, 2014), EF is associated with a diverse network of frontal regions. Contemporary reviews of lesion studies have not identified a one-to-one relationship between any EF task and specific regions (Alvarez & Emory, 2006). Executive deficits observed in patients may be the result of multiple attentional circuits connecting both frontal and posterior regions, which are adaptive to multiple contexts (Stuss & Alexander, 2007), some of which support a more focal relationship between individual regions and diverse EF functions. Although some studies can isolate focal regional associations using component analysis for specific tasks (Pa et al., 2010), generally the literature presents varied findings regarding the associations between D-KEFS tasks and discriminant regional specificity in the frontal lobes (Jurado & Rosselli, 2007). This provides evidence that substantial loss in the volume of prefrontal brain tissue results in lower EF. However, there is limited research that examines how regional volumes are related to individual components of EF.

The underlying factor structure of the D-KEFS tasks may have contributed to mixed neuroimaging findings. Although the D-KEFS tasks are designed to assess different aspects of EF functioning (Delis et al., 2001), investigations into the factor structure of the D-KEFS have found that many of the tasks load onto broad component factors associated with monitoring or inhibition (Latzman & Markon, 2010). Even these broad component factors, such as inhibition, have demonstrated very strong correlations with a common (“unitary”) EF component (Friedman, Miyake, Robinson, & Hewitt, 2011). This suggests that the variability among D-KEFS tasks within the same factor do not necessarily imply a separable profile of strengths and weaknesses. A unitary factor for D-KEFS tasks has been identified, as many EF tasks are shown to be saturated with a common factor, as well as a specific factor (Latzman & Markon, 2010).

Therefore, any given D-KEFS task score may result from either a specific component of EF, or instead, a common factor of EF. The pattern of neural correlates for D-KEFS tasks may support the construction of EF as a unitary (aspects of EF ability common to all tasks) and task (diverse EF skills) level, and guide the interpretability of D-KEFS score variation.

Using structural neuroimaging, we examined how the Tower, Verbal Fluency, Design Fluency, Color–Word Interference, and Trail Making Test tasks related to regional prefrontal volumes. We assessed how the “unitary” EF construct (average of D-KEFS scores) and “diverse” subcomponents of EF (individual D-KEFS task performance) were related to regional prefrontal brain volumes. The pericalcarine cortex from the occipital lobe, which is considered not strongly associated with EF tasks (Laird et al., 2005; Rottschy et al., 2012), was included as a control region to compare the strength of frontal relationships with “unitary” and “diverse” EF. Discriminant associations between these tasks and frontal brain regions would demonstrate construct validity for the D-KEFS tasks and elucidate the cortical organization of EF.

## Methods

### Participants

Anonymized data from the “Cross-Sectional Lifespan Connectomics Study” of the Enhanced Nathan Kline Institute – Rockland Sample (NKI-RS) were analyzed for the present study. NKI-RS is an open-access, cross-sectional community sample of persons from Rockland County, New York (Nooner et al., 2012). Ethnic and economic demographics of Rockland County closely resemble that of the United States (U.S. Census Bureau, 2010), increasing the generalizability of NKI-RS to the broader U.S. population. Zip code-based recruitment was conducted through mail, print advertising, and electronic advertising in Rockland County. Enrollment efforts prioritized representativeness of Rockland County to balance the proportion of key demographic variables (e.g., age, sex, race) in NKI-RS. Eligibility criteria were designed for inclusivity (Nooner et al., 2012). Nearly half of recruited individuals met criteria for at least one Diagnostic Statistical Manual - 4th Edition (DSM-IV, American Psychiatric Association, 2000) diagnosis based on a semi-structured clinical interview. Recruited adults were excluded from NKI-RS if screening indicated severe psychiatric disorders, severe developmental disorders, current suicidal or homicidal ideation, severe cerebral trauma, severe neurodegenerative disorders, substance dependency issues occurring in the previous two years, history of psychiatric hospitalization, current pregnancy, or magnetic resonance imaging (MRI) contraindications.

De-identified phenotypic and neuroimaging data from 645 adult “Cross-Sectional Lifespan Connectomics Study” NKI-RS participants was available at the start of this study (October, 2016). We selected adult participants (ages 20 to 85) to ensure adequate brain maturation (Dosenbach et al., 2010; Johnson, Blum, & Giedd, 2009) and decreased rate of late-life chronic illness (Wolff, Starfield, & Anderson, 2002). Eligibility for data analysis required complete MRI and D-KEFS data. Participants with incomplete or poor quality of structural brain images were excluded (*n* = 5). The final sample consisted of 478 participants (*M* = 48.88, *SD* = 17.43). Sixty-seven percent of the sample identified their sex as female. Participants were 76.15% white, 16.53% African American, and 3.97% Asian. With respect to ethnicity, 9.83% of participants identified as Hispanic. Nine percent of participants were left-handed. Average years of education were 15.64 (*SD* = 2.21, range = 9 – 24); 27.20% of participants had 18 or more years of education. All were proficient in English, though 5.65% of participants listed it as their second language.

### Procedures

All participants completed a medical evaluation, psychiatric interview, self-report questionnaires, a standardized battery of cognitive tests (including the D-KEFS), and neuroimaging at NKI. See Nooner et al. (2012)for a complete list of assessments and the participant assessment schedule. Participants enrolled in NKI-RS before September 15, 2015 engaged in a two-day research protocol. After that date, the participant schedule was abbreviated to a one-day protocol following an hour-long screening visit. The D-KEFS was administered by trained research assistants in accordance with the administration manual under the supervision of licensed neuropsychologists. Participants completed the D-KEFS on the same day as the MRI. Data use agreement was approved by NKI-RS and data handling procedures were accepted by the Institutional Review Board at Suffolk University.

### Measures

#### Executive function (EF)

The following five out of nine available tasks were used from the D-KEFS: Trail Making (TM), Design Fluency (DF), Color Word Interference (CWI), Tower, and Verbal Fluency (VF). These tasks were selected to broadly assess EF using classical paradigms from clinical neuropsychology. The five D-KEFS tasks were administered to both one-day and two-day NKI-RS participants, which maximized the sample size for study analyses. The primary performance measure for each task, described in greater detail below, was selected based on the literature (Delis et al., 2004, 2001; Strauss, Sherman, & Spreen, 2006).

##### Color–Word Interference (CWI)

The D-KEFS Color-Word Interference Test (CWI) builds upon a classic measure of cognitive control developed by Stroop (1935). It is intended to measure working memory, inhibitory control, cognitive flexibility, semantic activation, and response selection (Strauss et al., 2006). The task consists of four conditions; participants were required to name colored squares (1), read words indicating colors printed in black ink (2), name the incongruous ink color of printed color words (3), and switch between naming the ink color and reading the color words (4). This study only contains the results from Condition 4 (inhibition and switching), which requires participants to inhibit an overlearned response (i.e., naming color of the words while ignoring their semantic content) and demonstrate flexibility by set-shifting. Scoring is based on completion time, with faster times indicating better performance. According to Strauss et al. (2006), reliability for the CWI in the Delis et al. (2001)normative sample ranged from marginal (coefficients between .60 and .69) for test-retest and adequate for internal consistency (.70 to .79). Convergent validity for the CWI has been evidenced through significant associations with the Wisconsin Card Sorting Test for the constructs of inhibition (*r* = -.53) and inhibition and switching (*r* = -.31) (Delis et al., 2001), which is an established neuropsychological test of set-shifting (Heaton et al., 1993).

##### Tower

The D-KEFS Tower Test is designed to measure planning, problem solving, inhibition and the ability to resolve conflicts between goals and subgoals (Goel & Grafman, 1995; Morris, Miotto, Feigenbaum, Bullock, & Polkey, 1997;Strauss et al., 2006). It is comparable to other tower paradigms (e.g., Tower of London; Marchegiani, Giannelli, & Odetti, 2010) used by neuropsychologists to assess EF (Rabin et al., 2005). Participants are asked to move the five disks across the three pegs to construct a target tower using the fewest number of moves possible. This paradigm has two rules that place demands on planning capacity: (1) only one disk can be moved at a time and, (2) no disk may be placed on top of a smaller disk. The minimum number of moves required to solve each puzzle increases as participants progress through the nine items, which necessitates greater problem-solving and planning ability. Items were timed for discontinuation criteria. The towers that were successfully completed within the time limit were awarded one point, with additional points given for completing the tower using fewer moves (range = 0 to 4 points). The scores for each of the nine items were totaled to produce a total achievement score (range = 0 to 30 points). Delis et al. (2001)reported an overall moderate test-retest reliability for total achievement score (*r* = .40) in their normative sample, with the lowest value being 0.38 in ages 50-89. According to Strauss et al. (2006), the total achievement score had marginal internal consistency reliability across all age groups. Convergent validity for the D-KEFS Tower has been demonstrated through significant associations with subcomponents of the total achievement score on the Tower of London test including total move score (*r* = .47), total initiation time (*r* = .58), and total correct (*r* = .42) (Larochette, Benn, & Harrison, 2009).

##### Trail Making (TM)

The D-KEFS Trail Making Test (TM) is a modified version of the traditional trail making paradigm (Army Individual Test Battery, 1944; Reitan, 1955) which is one of the most widely used tests used by clinical neuropsychologists (Rabin et al., 2005). It is intended to assess set-shifting, attention, processing speed, and visuomotor scanning (Arbuthnott & Frank, 2000; Sánchez-Cubillo et al., 2009). Participants are asked to connect several targets (numbers, letters, dotted lines) that are scattered across the test form. The D-KEFS TM consists of five conditions: (1) visual scanning, (2) number sequencing, (3) letter sequencing, (4) number-letter switching, and (5) motor speed. When an error was committed by a participant, the tester instructed them to return to the last correct placement before proceeding. The primary performance measure was completion time, in seconds, for Condition 4. The trail making paradigm has demonstrated strong inter-rater reliability (*r* = .90) (Fals-Stewart, 1992). Test-retest reliability for TM conditions ranged from 0.38 (switching) to 0.77 (motor speed), with most coefficients within the moderate range for all age groups (Delis et al., 2001). D-KEFS TM number-letter switching (Condition 4) has demonstrated predictive validity via significant an association with functional status (*r* = .56) (Mitchell & Miller, 2008).

##### Verbal Fluency (VF)

The D-KEFS Verbal Fluency Test (VF) is intended to measure category fluency, set-shifting, and letter fluency (Swanson, 2005). It is analogous to other verbal phonemic fluency tests, such as Controlled Word Association Test (COWAT; Benton, Hamsher, & Sivan, 1983). Participants must name as many unique words as possible in 60 seconds for three different prompts: by letter (Condition 1), category (Condition 2), and category switching (Condition 3). The sum of correct responses for Condition 3, which required participants to switch between naming unique fruit and furniture items, served as the primary outcome measure for VF. In the Straus et al. (2001) development sample, Condition 3 demonstrated marginal internal consistency (.60 to .69) and low test-retest reliability (< .60). The verbal fluency paradigm has demonstrated a significant relationship with dysexecutive behavior such as confabulation, perseveration, and temporal sequencing (Burgess, Alderman, Evans, Emslie, &Wilson, 1998).

##### Design Fluency (DF)

Letter and verbal fluency tasks are analogous to design fluency tasks, and have inspired tasks such as the D-KEFS Design Fluency Test (DF) (Delis et al., 2001). These tasks are intended to assess planning, cognitive flexibility, and visual-motor processing (Strauss et al., 2006; Suchy, Kraybill, & Gidley Larson, 2010). The DF presents participants with rows of boxes that contain an array of dots. Participants are then instructed to draw as many unique geometric designs as possible within 60 seconds. The paradigm consists of three conditions: connecting filled dots (1), connecting unfilled dots (2), and alternating connections between filled and unfilled dots (3). Successful task performance relies on observing the rules and restrictions while not repeating the designs of previous drawings. The sum of correct designs within the 60 second time limit for Condition 3, which requires inhibition and set-shifting, served as the primary outcome measure. The DF has evidenced low test-retest reliability (.32 to .58) (Delis et al., 2001). However, the design fluency paradigm has demonstrated good to excellent inter-rater reliability (Carter, Shore, Harnadek, & Kubu, 1998; Harter, Hart, & Harter, 1999).

#### Neuroimaging

Structural images were acquired using a 3T Siemens Trio scanner (T1 MPRAGE, voxel size = 1.0 x 1.0 x 1.0 mm, 176 slices, echo time = 2.52 ms, repetition time = 1900 ms, field of view = 250 mm). MPRAGE data obtained from the NKI-RS dataset are available in their raw form. Complete details about the MRI measurement parameters can be located on the NKI-RS website for MPRAGE (http://fcon_1000.projects.nitrc.org/indi/enhanced/NKI_MPRAGE.pdf). MPRAGE data included in the NKI-RS dataset were obtained in their unprocessed form (Digital Imaging and Communications in Medicine; DICOM) and converted to the .mgz format.

Cortical reconstruction and volumetric segmentation of MPRAGE images were automatically reconstructed using Freesurfer (Version 5.3), a publicly available software package for analyzing and visualizing structural and functional neuroimaging data (http://surfer.nmr.mgh.harvard.edu/). Briefly, FreeSurfer image processing includes motion correction (Reuter, Rosas, & Fischl, 2010), removal of non-brain tissue (Ségonne et al., 2004), automated Talairach transformation and estimation of total intracranial volume (Buckner et al., 2004), intensity normalization, segmentation of white and gray matter structures (Fischl et al., 2004, 2002), tessellation of white/gray matter boundaries, topical surface correction (Fischl, Liu, & Dale, 2001; Ségonne, Pacheco, & Fischl, 2007), and surface deformation (Dale, Fischl, & Sereno, 1999; Fischl & Dale, 2000). Each image was mapped into standard morphological space (MNI305; Collins, 1994), and volumes (mm^3^) were generated for cortical regions based on theDesikan-Killiany Atlas (Desikan et al., 2006). A quality assurance protocol for surface reconstruction accuracy was performed as reported previously (Waters, Mace, Sawyer, & Gansler, 2017).

Segmented cortical gray matter volumes were generated by FreeSurfer for each hemisphere. All thirteen frontal regions of interest (ROI) in the Desikan-Killiany Atlas (Desikan et al., 2006) generated by FreeSurfer were extracted for study analyses (https://surfer.nmr.mgh.harvard.edu/fswiki/CorticalParcellation). Additionally, the pericalcarine cortex from the occipital lobe was included as a control region for comparing the association between frontal ROI and the D-KEFS. Descriptive statistics for ROI volumes, averaged across hemisphere, are presented in Supplementary Table 1.

### Statistical analyses

Statistical analysis was conducted using R 3.4.0 (R Core Team, 2017) in RStudio 1.0.143 (RStudio Team, 2016) (See Appendix for R packages). R code for all study analyses has been made available for reproducibility with open-access NKI-RS data (See Supplemental R Script).

#### Pre-analysis

Timed D-KEFS tasks (TM, CWI) were inverted so that positive scores indicated better performance for all EF measures. TM and CWI were also log transformed to meet assumptions of normality. ROI volumes were divided by participants’ total intracranial volume for normalization (Buckner et al., 2004; Chen et al., 2016). Descriptive statistics were used to report participant demographics and scores for D-KEFS tasks. ROI volumes and D-KEFS task scores were visualized in the *ggplot2* package to assess normality and identify outliers. Independent-sample t-tests and Pearson correlations were used to explore associations between the D-KEFS tasks and demographic factors.

#### Mixed-effects modeling

Mixed-effects modeling was conducted with the *lme4* package. Mixed-effects models are a flexible tool for modeling a wide variety of correlated data that use fixed and random effects. Mixed-effects models are ideal for high dimensional neuroimaging data because they can handle multiple measurements per subject (i.e., volume for each ROI). Advantages over traditional linear regression include: (1) enhanced statistical power due to repeated observations (2) robustness to missing data, and (3) management of heteroscedasticity and non-spherical error variance for either participants or study items (Baayen, Davidson, & Bates, 2008). Mixed-effects modeling was used to quantify which ROIs were most related to both individual EF scores (diverse construct) and the scores together (unitary construct). Prior to mixed-effects modeling, continuous variables (age, education, D-KEFS scores, ROI volumes) were standardized to z-scores (*M* = 0, *SD* = 1).

In a linear mixed-effects model predicting volume (the criterion), D-KEFS task (factor, 5 levels), D-KEFS score (continuous), ROI (factor, 14 levels) were entered as fixed effects. Intercepts for participants were added as a random effect. Age, sex, and education were entered as covariates due to their relationship with brain structure and EF (Lezak, 2012). Hemisphere was also included as a covariate (as opposed to an interaction term) to simplify interpretability and because we did not make predictions regarding EF lateralization. Omnibus statistics for the mixed-effects model were summarized using the *anova* function. The D-KEFS score (i.e., overall EF ability) x ROI interaction was used to assess unitary EF. The D-KEFS score x D-KEFS task x ROI interaction examined diverse EF (i.e., CWI, Tower, TM, VF, and DF as differential predictors of ROI volumes). Slopes for the EF-volume relationship for each ROI were estimated using the *lstrends* function in the *lsmeans* package. Two-way (D-KEFS score x ROI) and three-way (D-KEFS task x D-KEFS score x ROI) interactions were refit using 1,000 bootstrapped samples (with replacement) to create 95% confidence intervals (CI) for the fixed effect coefficients.

## Results

### Pre-analysis

Descriptive statistics for raw scores on the D-KEFS tasks are presented in Table 1. TM (skewness = 2.01) and CWI (skewness = 1.40) were log transformed to meet assumptions of normality. Age was significantly associated with poorer performance on the CWI (*r* = −0.34, *p* < 0.001), TM (*r* = −0.29, *p* < 0.001), DF (*r* = −0.24, *p* < 0.001), Tower (*r* = −0.14, *p* < 0.01), and higher performance on VF (*r* = 0.12, *p* < 0.01). Education was significantly associated with better performance on VF (*r* = 0.28, *p* < 0.001), TM (*r* = 0.16, *p* < 0.001), Tower (*r* = 0.12, *p* < 0.01), DF (*r* = 0.10, *p* = 0.03) but not CWI (*r* = 0.06, *p* = 0.19). D-KEFS scores were not significantly associated with sex (*ps* > 0.05). Correlations between the D-KEFS tasks were significant (all *p*s < 0.001), with small (Tower and CWI, *r* = 0.21) to moderate (CWI and TM, *r* = 0.50) associations (see Table 3).

**Table 1.** Descriptive statistics and correlations for D-KEFS tasks.

| *Task* | *Descriptive statistics* |  |  |  |  |  | *Pearson correlations* |  |  |  |
| --- | --- | --- | --- | --- | --- | --- | --- | --- | --- | --- |
|  | *M* | *SD* | *Min* | *Max* | *Skew* | *Kurtosis* | *1* | *2* | *3* | *4* |
| 1. CWI | 55.42 | 14.79 | 29 | 127 | 1.4 | 3.44 |  |  |  |  |
| 2. TM | 81.69 | 38.94 | 25 | 240 | 2.01 | 4.84 | 0.47 |  |  |  |
| 3. DF | 8.04 | 2.66 | 0 | 16 | 0 | 0.11 | -0.43 | -0.44 |  |  |
| 4. VF | 40.97 | 12.11 | 9 | 77 | 0.19 | -0.05 | -0.28 | -0.36 | 0.23 |  |
| 5. Tower | 15.98 | 4.15 | 2 | 27 | -0.52 | 0.58 | -0.23 | -0.39 | 0.33 | 0.19 |

*Note.* Color Word Interference (CWI), Trail Making (TM), Design Fluency (DF), Tower, and Verbal Fluency (VF). All correlations were significant ( $ps < 0.001$ ).

**Table 2.** Summary of the mixed-effects model.

| *Predictor* | *Sum sq* | *Mean sq* | *Num df* | *Den df* | *F-value* | *P-value* |
| --- | --- | --- | --- | --- | --- | --- |
| task | 0.00 | 0.00 | 4 | 66163 | 0.00 | 1.00 |
| score | 0.05 | 0.05 | 1 | 66210 | 0.07 | 0.79 |
| region | 0.00 | 0.00 | 13 | 66163 | 0.00 | 1.00 |
| hemisphere | 0.00 | 0.00 | 1 | 66163 | 0.00 | 1.00 |
| age | 43.07 | 43.07 | 1 | 476 | 62.17 | < 0.001 |
| sex | 2.57 | 2.57 | 1 | 474 | 3.70 | 0.05 |
| education | 0.00 | 0.00 | 1 | 475 | 0.01 | 0.94 |
| task x score | 0.00 | 0.00 | 4 | 66445 | 0.00 | 1.00 |
| task x region | 0.00 | 0.00 | 52 | 66163 | 0.00 | 1.00 |
| **score x region (unitary)** | **47.14** | **3.63** | **13** | **66163** | **5.234** | **< 0.001** |
| **task x score x region (diverse)** | **39.50** | **0.76** | **52** | **66163** | **1.096** | **0.29** |

*Note.* Unitary and diverse EF were bolded. Sum of squares (Sum sq); mean square (Mean sq); numerator (Num df) and denominator (Den df) degrees of freedom.

**Table 3.** Bootstrapped estimates for the D-KEFS score x ROI interaction predicting volume.

| *ROI* | *Slope* | *Lower CI* | *Upper CI* |
| --- | --- | --- | --- |
| **superior frontal** | **0.051** | **0.016** | **0.087** |
| **rostral middle frontal** | **0.047** | **0.005** | **0.085** |
| **lateral orbitofrontal** | **0.040** | **0.005** | **0.075** |
| medial orbitofrontal | 0.003 | -0.034 | 0.041 |
| paracentral | 0.001 | -0.040 | 0.038 |
| caudal middle frontal | -0.002 | -0.044 | 0.041 |
| caudal anterior cingulate | -0.002 | -0.048 | 0.044 |
| pars opercularis | -0.002 | -0.040 | 0.038 |
| precentral | -0.007 | -0.042 | 0.029 |
| pars triangularis | -0.009 | -0.047 | 0.031 |
| rostral anterior cingulate | -0.014 | -0.049 | 0.021 |
| pericalcarine | -0.017 | -0.077 | 0.042 |
| pars orbitalis | -0.033 | -0.075 | 0.010 |
| **frontal pole** | **-0.044** | **-0.087** | **-0.001** |

*Note.* Bootstrapped ( $B = 1,000$ ) slopes and 95% confidence intervals (CI) from the D-KEFS score x region of interest (ROI) interaction. Slope values are the average of D-KEFS scores over volumes (z-standardized). Significant levels ( $p < 0.05$ ) were bolded.

### Mixed effects modeling

Results from the mixed-effects model are reported in Table 2. Age significantly predicted frontal lobe volume (*p* < 0.001). Every 1 *SD* increase in age (17.45 years) was associated with a 0.20 *SD* decrease (95% CI: −0.24, −0.15) in volume. Education (*B* = 0.002; 95% CI: −0.047, 0.051) and sex (female, *B* = .03; 95% CI: −0.03, 0.09) did not significantly predict frontal lobe volume. D-KEFS score, which represented participants’ overall EF ability, did not significantly predict frontal lobe volume (*B* = 0.0002; 95% CI: −0.0003, 0.0008). (Table 2 here)

#### Unitary EF

As presented in Table 2, the two-way D-KEFS score x ROI interaction was significant (*p* < 0.001). Table 3 presents bootstrapped slopes and 95% CI for all levels of the D-KEFS score x ROI interaction, which are visualized as horizontal error bars in Figure 1. Unitary EF (i.e., without the task interaction term) significantly predicted volume in four frontal ROIs. Every 1 *SD* increase in overall EF ability was significantly associated with 0.051, 0.047, and 0.040 *SD greater* volume in the superior frontal gyrus (95% CI: 0.016, 0.087), rostral middle frontal gyrus (95% CI: 0.005, 0.085), and lateral orbitofrontal cortex (95% CI: 0.005, 0.075), respectively. Conversely, every 1 *SD* increase in overall EF ability was significantly associated with 0.044 *SD lower* volume in the frontal pole (95% CI: −0.087, −0.001). (Table 3 and Figure 1 here)

**Figure 1.**
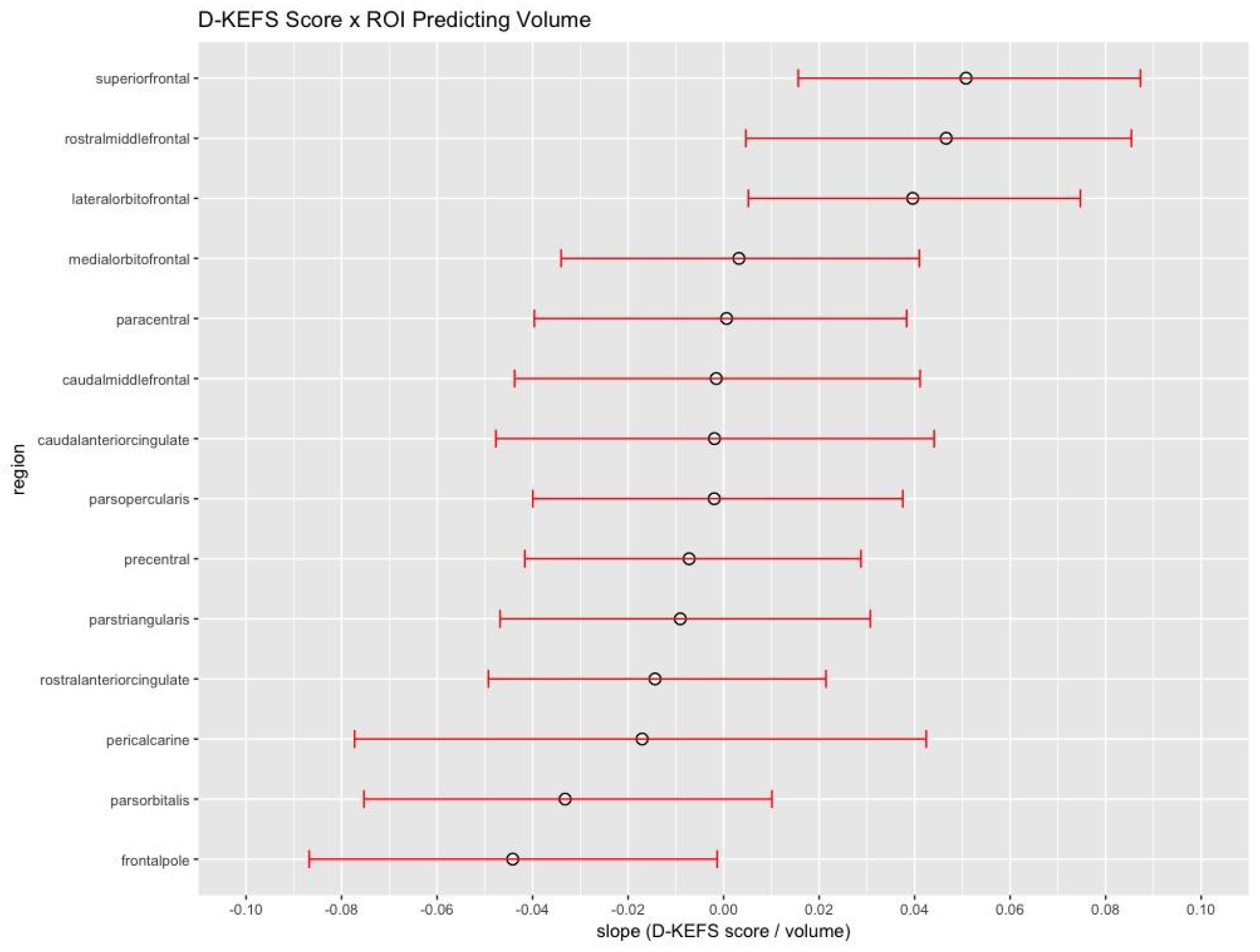
Visualization of the bootstrapped slopes (black circle) and 95% confidence intervals (error bars) from Table 3. D-KEFS scores and volumes were z-standardized.

#### Diverse EF

As presented in Table 2, the three-way D-KEFS task x D-KEFS score x ROI interaction was not significant (*p* = 0.29). Table 4 lists the 7 levels from the bootstrapped three-way interaction that significantly predicted ROI volume. Supplementary Figure 1 displays the results from Table 4 on the left hemisphere of FreeSurfer’s inflated brain template. Every 1 *SD* increase in DF performance was significantly associated with a 0.103 *SD greater* volume in the lateral orbitofrontal cortex (95% CI: 0.047, 0.155). The TM (*B* = 0.062; 95% CI: 0.006, 0.121) significantly predicted larger rostral middle frontal gyrus volume. The CWI (*B* = 0.099; 95% CI: 0.047, 0.155), TM (*B* = 0.073; 95% CI: 0.017, 0.131), and DF (*B* = 0.053; 95% CI: 0.002, 0.101) significantly predicted larger superior frontal gyrus volume. Conversely, higher performance on DF (*B* = −0.075; 95% CI: −0.143, −0.011) and CWI (*B* = −0.066; 95% CI: −0.128, −0.006) significantly predicted lower frontal pole volume. Bootstrapped slopes and 95% CI for all 70 levels of the three-way interaction are reported in Supplementary Table 2. (Table 4 here)

**Table 4.** Bootstrapped estimates for D-KEFS score x D-KEFS task x ROI interaction predicting volume.

| *Task* | *ROI* | *Slope* | *Lower CI* | *Upper CI* |
| --- | --- | --- | --- | --- |
| DF | lateral orbitofrontal | 0.103 | 0.047 | 0.155 |
|  | superior frontal | 0.053 | 0.002 | 0.102 |
|  | frontal pole | -0.075 | -0.143 | -0.011 |
| CWI | superior frontal | 0.099 | 0.047 | 0.155 |
|  | frontal pole | -0.066 | -0.128 | -0.006 |
| TM | superior frontal | 0.073 | 0.017 | 0.131 |
|  | rostral middle frontal | 0.062 | 0.006 | 0.121 |

*Note.* Bootstrapped ( $B = 1,000$ ) slopes and 95% confidence intervals (CI) from the D-KEFS
score x D-KEFS task x region of interest (ROI) interaction. Slope values are individual D-KEFS scores over volumes (z-standardized). Only statistically significant levels ( $p < 0.05$ ) are shown;
Supplementary Table 2 shows all levels.

## Discussion

The neural correlates found for the D-KEFS in this study support both the unitary (average of D-KEFS scores) and diverse (individual D-KEFS scores) prefrontal modularity of EF. Our findings are consistent with the evidence that brain volume is related to neuropsychological ability, but also further distinguishes how the brain and behavior are related (Cipolotti & Warrington, 1995). The average of D-KEFS scores were more broadly associated with prefrontal volume, whereas individual D-KEFS scores were more narrowly but strongly related to specific ROIs. Unitary EF was significantly associated with four of 13 frontal ROIs and the magnitude of the relationships (| *z*-score slope |) ranged from 0.040 to 0.051 (see Table 3 and Figure 1. Diverse EF skills were differentially associated with two or three frontal ROIs, respectively (see Table 4), but to a stronger extent (range | *z* | = 0.053 to 0.103). This aligns with evidence from factor analytic and neuroimaging studies that have identified both unitary and diverse contributions to the functional and cortical organization of EF (Elliott, 2003; Huizinga, Dolan, & van der Molen, 2006; Jurado & Rosselli, 2007; Miyake et al., 2000). Mixed-effects modeling of D-KEFS and structural neuroimaging data with bootstrap resampling offered a robust method for simultaneously estimating the hierarchical relationship between EF and the prefrontal cortex. The likely negligible relationship of EF with the occipital control region provided further validation of the observed and expected EF-frontal associations. Perhaps the unitary versus diverse debate of EF is fueled by previous studies’ limitations when incorporating the relative contributions of both aspects (Miyake et al., 2000).

Unitary EF significantly and positively predicted volume in the superior frontal, rostral middle frontal, and lateral orbitofrontal gyri. This confirms the relationship between larger lateral prefrontal cortex and better EF performance found for healthy adults in the Yuan and Raz (2014)meta-analysis. The D-KEFS tasks used for unitary EF may recruit multiple lateral prefrontal cortex regions involved in broad component functions of monitoring or inhibition (Friedman et al., 2011; Latzman & Markon, 2010). These metacognitive or cognitive control skills, which are routinely assessed on neuropsychological tests, have been localized to the lateral prefrontal cortex (Ardila, 2013). Nevertheless, our findings contribute to emerging evidence that aggregate measurements of EF may serve broader but less robust frontal neural correlates than diverse EF skills (Waters et al., 2017). Unitary EF in this study is more comparable to EF aggregates, such as the sum of the six Functional Assessment Battery (Dubois, Slachevsky, Litvan, & Pillon, 2000) tasks, than factor analytic composite of unitary EF (Gansler et al., 2015; Miyake & Friedman, 2012). Future research should examine the importance of factor analytic extraction, which might be advantageous for handling task impurity and measurement error (Friedman et al., 2008), as it compares to these aggregate measures.

Mixed-effects modeling revealed dissociable relationships between diverse EF and frontal ROIs, the strength of which varied by D-KEFS task. The strongest signal, between DF and the lateral orbitofrontal cortex, may reflect the prevention of repetition errors (Possin et al., 2012) through self-monitoring and subsequent behavior modification based on a series of contingencies (correct designs are rewarding, repeating designs is punishing) (Bechara, 2004; Kringelbach & Rolls, 2004; O’Doherty, Kringelbach, Rolls, Hornak, & Andrews, 2001), as well as responsivity to social cues provided by the test administrator (Liu et al., 2007). The significant correlation between the rostral middle frontal gyrus aligns with the role of the dorsolateral prefrontal cortex in cognitive flexibility (Kopp et al., 2015; Stuss et al., 2001; Varjacic, Mantini, Demeyere, & Gillebert, 2018). The overlapping relationship between DF, CWI, and TM with the superior frontal gyrus (see Figure 1) could be attributed to the region’s role in working memory and cognitive flexibility (Boisgueheneuc et al., 2006; D’Esposito, 2007; Müller & Knight, 2006; Wager & Smith, 2003). These broad component cognitive functions are evident in the design of the D-KEFS DF, CWI, and TM tasks. Delis et al. (2001)created each D-KEFS condition to isolate critical cognitive functions (e.g., cognitive flexibility on TM Condition 4) to increase sensitivity to mild brain damage (Strauss et al., 2006). Parsing out response properties (e.g., motor speed on the classic Trail Making Test), may have instead resulted in homogeneity among D-KEFs tasks that drive EF-frontal saturation.

Our findings did not uniformly support the “bigger is better” hypothesis of brain-behavior relationships in non-patients (Yuan & Raz, 2014). Interestingly, better performance on both unitary and diverse (DF, CWI) measurements of EF were significantly associated with smaller frontal pole volume. Previous research suggests that the frontal pole is specialized for disengaging cognitive control from the task at hand (Mansouri, Buckley, Mahboubi, & Tanaka, 2015) and exploring the value of external tasks (Burgess, Gilbert, & Dumontheil, 2007; Ramnani & Owen, 2004) (Burgess, Golbert, & Dumontheil, 2007; Ramnani & Owen, 2004)—cognitive processes that may be contraindicated during the highly structured D-KEFS testing environment. Further research is needed to confirm the negative EF-frontal pole relationship in this study, particularly given that its upper-bound confidence interval approached non-significance (see Figure 1). Functional neuroimaging research is better suited to compare the frontal pole correlation of D-KEFS tasks that involve and do not involve rapid disengagement, such as CWIT Conditions 3 and 4, respectively.

Tower and VF scores were not significantly related to frontal volumes. Perhaps relationships in those regions were small, vary substantially between people, or stronger relationships could exist with other brain regions. Category (as opposed to letter) fluency may be mediated by the temporal cortex (Baldo, Schwartz, Wilkins, & Dronkers, 2006) and Tower performance may be distributed across a fronto-parietal network (Guevara, Rizo Martínez, Robles Aguirre, & Hernández González, 2012; Lazeron et al., 2000).

The results that indicated relationships of prefrontal volumes with both unitary and diverse EF confers clinical implications for neuropsychologists. We observed weak correlates with unitary EF, and more diffuse damage is related to larger EF deficits in lesion studies (Stuss & Alexander, 2000). In conjunction with our findings, a composite of EF scores would be suitable for assessing more diffuse frontal lobe damage, such as hypoxia or secondary brain injury after head trauma. Another clinical application for the composite score is the assessment of dysexecutive syndrome (Baddeley & Wilson, 1988), a confluence of EF and behavioral deficits from prefrontal cortex dysfunction (Wilson, Evans, Emslie, Alderman, & Burgess, 1998). Behavioral disturbances in dysexecutive syndrome (e.g., apathy, disinhibition) are broadly associated with the prefrontal cortex, including dorsolateral and orbitofrontal regions (Duffy & Campbell, 2001; Gansler, Huey, Pan, Wasserman, & Grafman, 2017; Lichter & Cummings, 2001). Executive deficits and dysexecutive behavior predict loss of independence (Godefroy et al., 2010) and psychosocial intervention outcomes (Beaudreau, Rideaux, O’Hara, & Arean, 2015;Goodkind et al., 2015), which underscores the need for accurate neuropsychological assessment of frontal systems. When localized brain injuries are suspected, such as a tumor or penetrating head injury, separable EF measures could be emphasized. Based on the robust estimation in this study, DF, CWI, and TM can discriminate functions of orbitofrontal, superior frontal, and rostral middle frontal regions, respectively.

Several study limitations warrant discussion. First, one condition from each task was selected to represent classical EF paradigms from neuropsychology, which precluded analyses of the range of process variables made available by the D-KEFS. Future studies could use penalized regression methods, such as LASSO (least absolute shrinkage and selection operator), to select optimal EF predictors of frontal lobe structure. While this study assessed five neuropsychological paradigms of EF, it was by no means an exhaustive list (see Table 8-1 in Strauss et al., 2006) and some D-KEFS tasks (Sorting Test) were excluded because they were administered to fewer NKI-RS participants. Moreover, the reported brain-behavior relationships could vary between D-KEFS and other EF tasks. It is important to note that the study is cross-sectional; therefore, observations are correlational and causality in the EF-frontal relationships cannot be inferred. Although NKI-RS is demographically and socioeconomically generalizable to the US, the contribution of this study to clinical neuropsychology is predicated on such brain-behavior relationships operating in both clinical and non-clinical samples. In accordance with the validation of the D-KEFS in lesion studies, the modest effect sizes for EF and frontal volume found in this study are expected to be stronger in neuropsychiatric populations (Waters, Swenson, & Gansler, 2018).

Discovery of brain-behavior relationships requires simultaneous consideration of the multiple levels across which higher cognitive function operates. Research that separates task-general variance in neurocognitive domains from task-specific variance can further evaluate testing paradigms as indices of brain integrity. Our estimation of the association between EF paradigms from clinical neuropsychology and prefrontal cortex structure, at two levels, serves as critical benchmarks as the field migrates toward the use of computerized neuroscience tasks (Computerized Neurocognitive Battery,Gur, 2001; NIH toolbox,Gershon et al., 2010) for this same purpose. The smaller effect sizes in this study underscore the importance of leveraging larger neuroimaging datasets to estimate population parameters of neurocognitive function. Indeed, meta-analytic reports of EF frontal neural correlates may be inflated by overfitting small samples sizes (Levine, Asada, & Carpenter, 2009; Zhang, Xu, & Ni, 2013). We expect more conservative estimates that are consistent with our results as the field moves toward larger, multi-modal datasets of EF (Pan et al., in press).

## Acknowledgement

The authors thank the NKI-Rockland Sample Initiative for collecting and providing the data used in these analyses.

